# Novel *Wolbachia* strains in *Anopheles* malaria vectors from Sub-Saharan Africa

**DOI:** 10.1101/338434

**Authors:** Claire L Jeffries, Gena G Lawrence, George Golovko, Mojca Kristan, James Orsborne, Kirstin Spence, Eliot Hurn, Janvier Bandibabone, Michael L Tantely, Fara N Raharimalala, Kalil Keita, Denka Camara, Yaya Barry, Francis Wat’senga, Emile Z Manzambi, Yaw A Afrane, Abdul R Mohammed, Tarekegn A. Abeku, Shivanand Hegde, Kamil Khanipov, Maria Pimenova, Yuriy Fofanov, Sébastien Boyer, Seth R Irish, Grant L Hughes, Thomas Walker

## Abstract

*Anopheles (An.)* mosquitoes contain bacteria that can influence *Plasmodium* parasites. *Wolbachia*, a common insect endosymbiont, has historically been considered absent from *Anopheles* but has recently been found in *An. gambiae* populations. Here, we assessed a range of *Anopheles* species from five malaria-endemic countries for *Wolbachia* and *Plasmodium* infection. Strikingly, we found *Wolbachia* infections in *An. coluzzii, An. gambiae* s.s, *An. arabiensis, An. moucheti* and *An.* species ‘A’ increasing the number of *Anopheles* species known to be naturally infected by this endosymbiont. Molecular analysis suggests the presence of phylogenetically diverse novel strains, while qPCR and 16S rRNA sequencing indicates that *Wolbachia* is the dominant member of the microbiota in *An. moucheti* and *An.* species ‘A’. We found no evidence of *Wolbachia/Asaia* co-infections, and presence of these endosymbionts did not have significant effects on malaria prevalence. We discuss the importance of novel *Wolbachia* strains in *Anopheles* and potential implications for disease control.

## Introduction

Malaria is transmitted to humans through inoculation of *Plasmodium (P.)* sporozoites during the infectious bite of an infected female *Anopheles (An.)* mosquito. The genus *Anopheles* consists of 475 formally recognised species with ~40 vector species/species complexes responsible for the transmission of malaria at a level of public health concern [1]. During the mosquito infection cycle, *Plasmodium* parasites encounter a variety of resident microbiota both in the mosquito midgut and other tissues. Numerous studies have shown that certain species of bacteria can inhibit *Plasmodium* development [2–4]. For example, *Enterobacter* bacteria that reside in the *Anopheles* midgut can inhibit the development of *Plasmodium* parasites prior to their invasion of the midgut epithelium [5,6]. *Wolbachia* endosymbiotic bacteria are estimated to naturally infect ~40% of insect species [7] including mosquito vector species that are responsible for transmission of human diseases such as *Culex (Cx.) quinquefasciatus* [8–10] and *Aedes (Ae.) albopictus* [11,12]. Although *Wolbachia* strains have been shown to have variable effects on arboviral infections in their native mosquito hosts [13–15], transinfected *Wolbachia* strains have been considered for mosquito biocontrol strategies, due to a variety of synergistic phenotypic effects. Transinfected strains in *Ae. aegypti* and *Ae. albopictus* provide strong inhibitory effects on arboviruses, with maternal transmission and cytoplasmic incompatibility enabling introduced strains to spread through populations [16–22]. Open releases of *Wolbachia*-transinfected *Ae. aegypti* populations have demonstrated the ability of the *w*Mel *Wolbachia* strain to invade wild populations [23] and provide strong inhibitory effects on viruses from field populations [24], with releases currently occurring in arbovirus endemic countries such as Indonesia, Vietnam, Brazil and Colombia (https://www.worldmosquitoprogram.org).

The prevalence of *Wolbachia* in *Anopheles* species has not been extensively studied, with most studies focused in Asia using classical PCR-based screening, and up until 2014 there has been no evidence of resident strains in mosquitoes from this genus [25–29]. Furthermore, significant efforts to establish artificially-infected lines were, up until recently, also unsuccessful [30]. Somatic, transient infections of the *Wolbachia* strains *w*MelPop and *w*AlbB in *An. gambiae* were shown to significantly inhibit *P. falciparum* [31] but the interference phenotype is variable with other *Wolbachia* strain-parasite combinations [32–34]. A stable line was established in *An. stephensi*, a vector of malaria in southern Asia, using the *w*AlbB strain and this was also shown to confer resistance to *P. falciparum* infection [35]. One potential reason postulated for the absence of *Wolbachia* in *Anopheles* species was thought to be due to the presence of other endosymbiotic bacteria, particularly from the genus *Asaia* [36]. This acetic acid bacterium is stably associated with several *Anopheles* species and is often the dominant species in the mosquito microbiota [37]. In laboratory studies, *Asaia* has been shown to impede the vertical transmission of *Wolbachia* in *Anopheles* [36] and was shown to have a negative correlation with *Wolbachia* in mosquito reproductive tissues [38].

Recently, resident *Wolbachia* strains have been discovered in the *An. gambiae* s.l. complex, which consists of multiple morphologically indistinguishable species including several major malaria vector species. *Wolbachia* strains (collectively named *w*Anga) were found in *An. gambiae* s.l. populations in Burkina Faso [39] and Mali [40], suggesting that *Wolbachia* may be more abundant in the *An. gambiae* complex across Sub-Saharan Africa. Globally, there is a large variety of *Anopheles* vector species (~70) that have the capacity to transmit malaria [41] and could potentially contain resident *Wolbachia* strains. Additionally, this number of malaria vector species may be an underestimate given that recent studies using molecular barcoding have also revealed a larger diversity of *Anopheles* species than would have be identified using morphological identification alone [42,43].

In this study, we collected *Anopheles* mosquitoes from five malaria-endemic countries; Ghana, Democratic Republic of the Congo (DRC), Guinea, Uganda and Madagascar, from 2013-2017. Wild-caught adult female *Anopheles* were screened for *P. falciparum* malaria parasites, *Wolbachia* and *Asaia* bacteria. In total, we analysed mosquitoes from 17 *Anopheles* species that are known malaria vectors or implicated in transmission, and some unidentified species, discovering five species of *Anopheles* with resident *Wolbachia* strains; *An. coluzzii* from Ghana, *An. gambiae* s.s., *An. arabiensis, An. moucheti* and *Anopheles* species ‘A’ from DRC. Using *Wolbachia* gene sequencing we show that the resident strains in these malaria vectors are diverse, novel strains and qPCR and 16S rRNA amplicon sequencing data suggests that the strains in *An. moucheti* and *An.* species ‘A’ are higher density infections, compared to the strains found in the *An. gambiae* s.l. complex. We found no evidence for either *Wolbachia-Asaia* co-infections, or for either endosymbiont having any significant effect on the prevalence of malaria in wild mosquito populations.

## Results

### Mosquito species and resident *Wolbachia* strains

*Anopheles* species composition varied depending on country and mosquito collection sites (**Table 1**). We detected *Wolbachia* in *An. coluzzii* (previously named M molecular form) mosquitoes from Ghana (prevalence of 4% - termed *w*Anga-Ghana) and *An. gambiae* s.s. (previously named S molecular form) from all six collection sites in DRC (prevalence range of 8-24%) in addition to a single infected *An. arabiensis* from Kalemie in DRC (**Figure 1, Table 1**). The molecular phylogeny of the ITS2 gene of *Anopheles gambiae* s.l. complex individuals (including both *Wolbachia*-infected and uninfected individuals analysed in our study) confirmed molecular species identifications made using species-specific PCR assays (**Figure 2**). Novel resident *Wolbachia* infections were detected in two additional *Anopheles* species from DRC; *An. moucheti* (termed *w*AnM) from Mikalayi, and *An.* species A (termed *w*AnsA) from Katana. Additionally, we screened adult female mosquitoes of *An.* species A (collected as larvae and adults) from Lwiro, a village near Katana in DRC, and detected *Wolbachia* in 30/33 (91%), indicating this resident *w*AnsA strain has a high infection prevalence in populations in this region. The molecular phylogeny of the ITS2 gene revealed *Wolbachia*-infected individuals from Lwiro and Katana are the same *An.* species A (**Figure 3**) previously collected in Eastern Zambia [43] and Western Kenya [44]. All ITS2 sequences were deposited in GenBank (accession numbers MH598414 – MH598445) **(Supplementary Table 1)**.

**Table 1.**
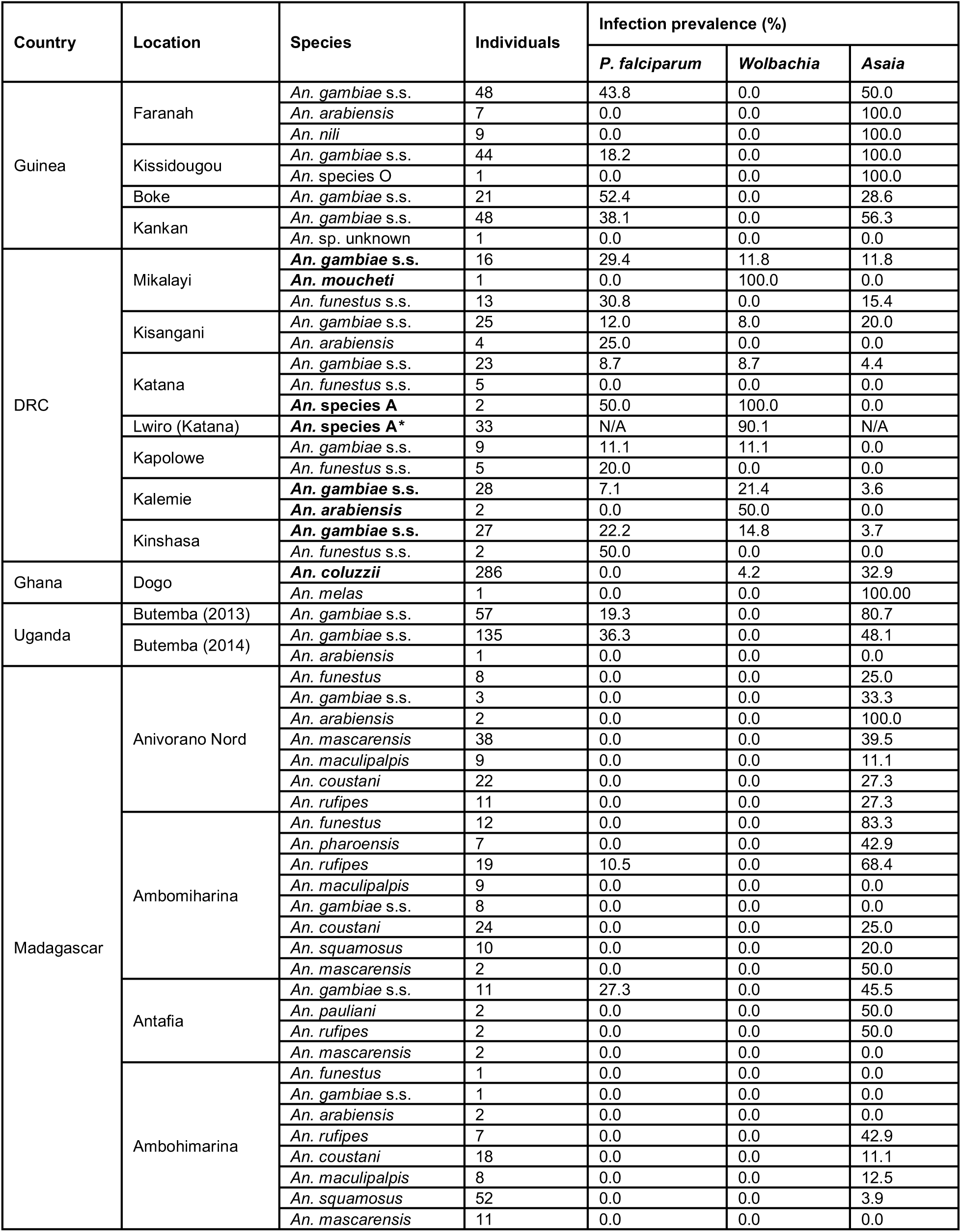
*Anopheles* mosquito species collected from locations within five malaria-endemic countries and *P. falciparum, Wolbachia* and *Asaia* prevalence rates. Species in different locations infected with *Wolbachia* are in bold. *Adult individuals from Lwiro (Katana), DRC were collected as both larvae and adults so have been excluded from *P. falciparum* and *Asaia* prevalence analysis.

**Figure 1.**
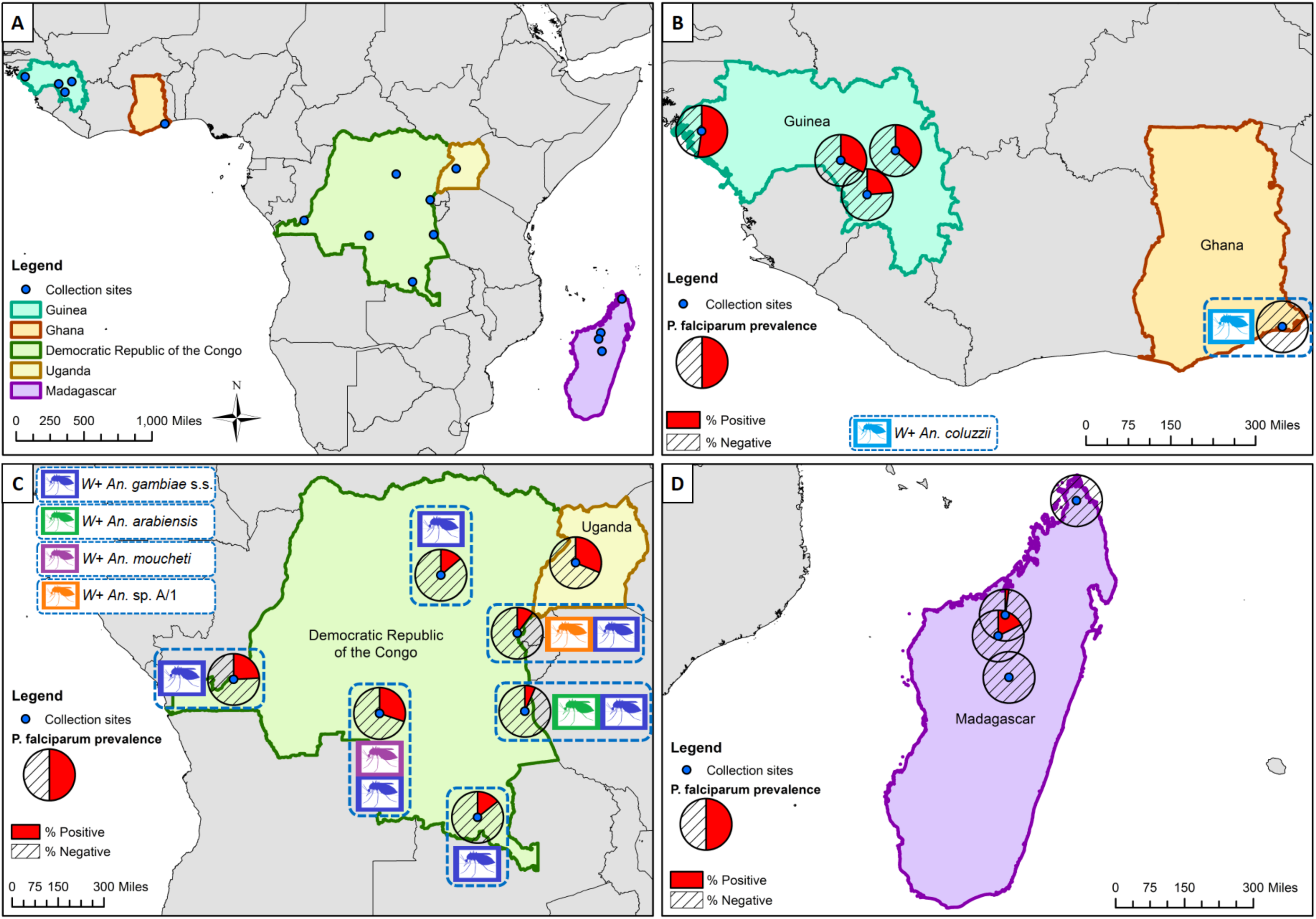
Locations of *Anopheles* species collections (including *Wolbachia*-infected species) and *P. falciparum* malaria prevalence rates in mosquitoes (across all species for each location). **A)** Overall map showing the five malaria-endemic countries where mosquito collections were undertaken. **B)** High malaria prevalence rates in Guinea, and *Wolbachia*-infected *An. coluzzii* from Ghana (no *P. falciparum* detected). **C)** *Wolbachia* strains in *An. gambiae* s.s., *An. arabiensis, An. species A* and *An. moucheti* from DRC and variable *P. falciparum* prevalence rate in DRC and Uganda. **D)** Low *P. falciparum* infection rates in Madagascar and no evidence of resident *Wolbachia* strains. (*W*+; *Wolbachia* detected in this species).

**Figure 2.**
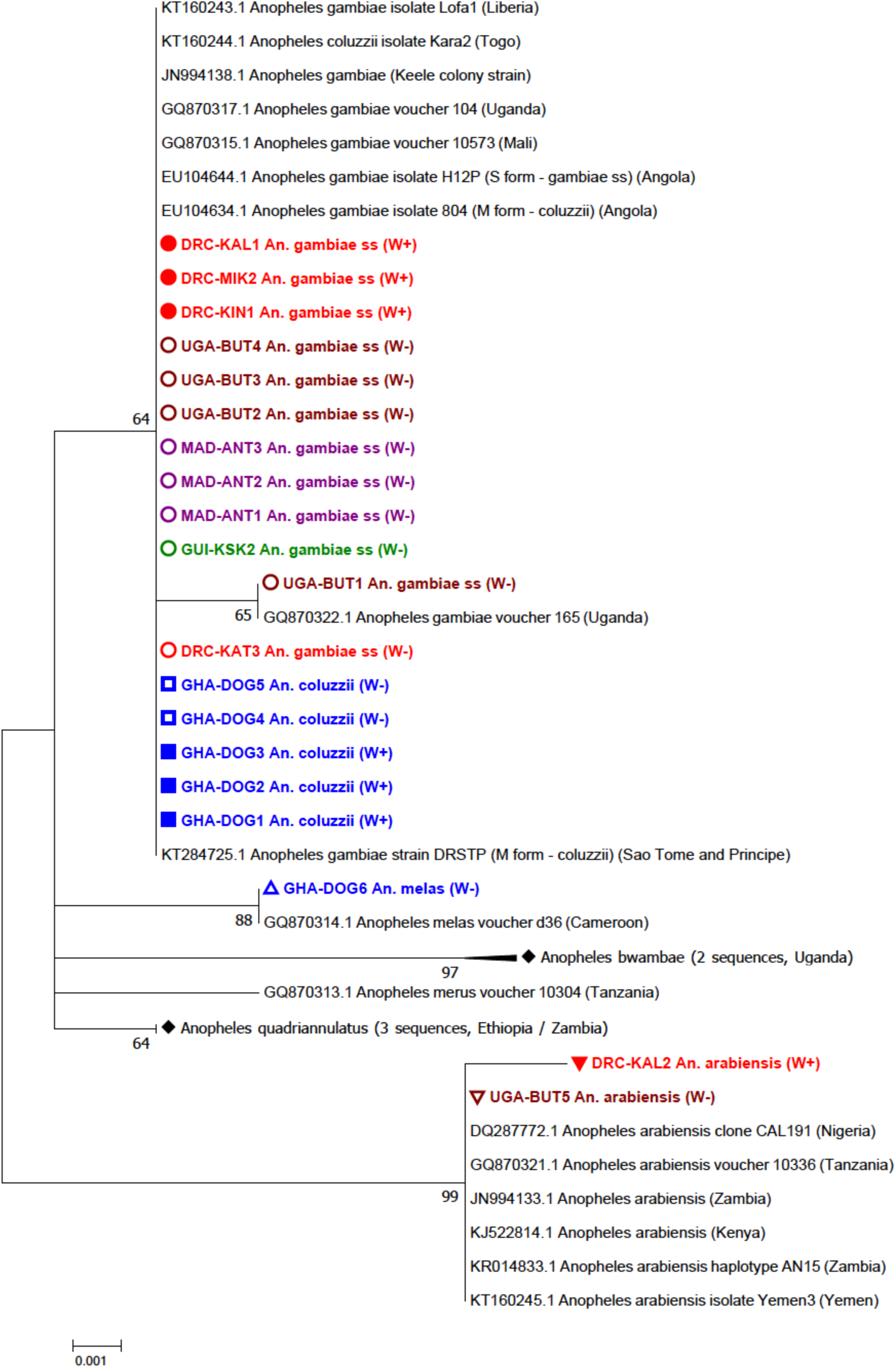
Maximum Likelihood molecular phylogenetic analysis of *Anopheles gambiae complex* ITS2 sequences from field-collected mosquitoes. The tree with the highest log likelihood (-785.65) is shown. The tree is drawn to scale, with branch lengths measured in the number of substitutions per site. The analysis involved 42 nucleotide sequences. There were a total of 475 positions in the final dataset. DRC = Democratic Republic of the Congo (red): KAL = Kalemie, MIK = Mikalayi, KIN = Kinshasa, KAT = Katana. GHA = Ghana (blue): DOG = Dogo. GUI = Guinea (green): KSK = Kissidougou. MAD = Madagascar (purple): ANT = Antafia. UGA = Uganda (maroon): BUT = Butemba. (W+; individual was *Wolbachia* positive, W-; individual was *Wolbachia* negative).

**Figure 3.**
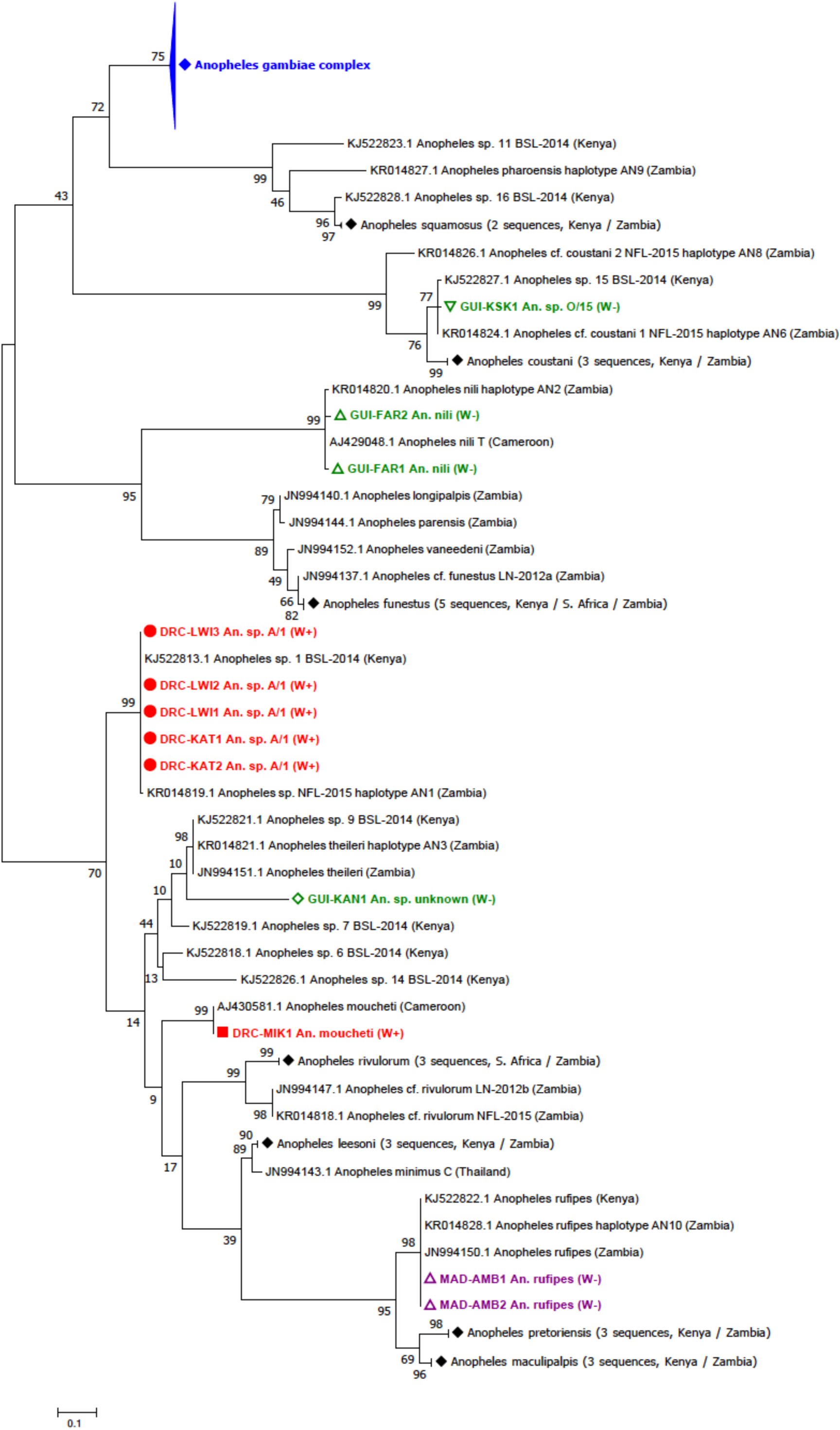
Maximum Likelihood molecular phylogenetic analysis of *Anopheles* ITS2 sequences from field-collected mosquitoes outside of the *An. gambiae* s.l. complex. The tree with the highest log likelihood (-3084.12) is shown. The tree is drawn to scale, with branch lengths measured in the number of substitutions per site. The analysis involved 118 nucleotide sequences. There were a total of 156 positions in the final dataset. DRC = Democratic Republic of the Congo (red): KAT = Katana, LWI = Lwiro, MIK = Mikalayi. GUI = Guinea (green): FAR = Faranah, KAN = Kankan, KSK = Kissidougou. MAD = Madagascar (purple): AMB = Ambomiharina. (W+; individual was *Wolbachia* positive, W-; individual was *Wolbachia* negative).

### *Wolbachia* strain typing

Phylogenetic analysis of the 16S rRNA gene demonstrated that the 16S sequences for these strains cluster with other Supergroup B strains such as *w*Pip (99-100% nucleotide identity) (**Figure 4a**). When compared to the resident *Wolbachia* strains in *An. gambiae* s.l. populations from Mali [40] and Burkina Faso [39], *w*Anga-Ghana is more closely related to the Supergroup B strain of *w*Anga from Burkina Faso. Although a resident strain was detected in *An. gambiae* s.s. and a single *An. arabiensis* from DRC through amplification of 16S rRNA fragments using two independent PCR assays [40,45], we were unable to obtain 16S sequences of sufficient quality to allow further analysis. The *Wolbachia* surface protein (wsp) gene has been evolving at a faster rate and provides more informative strain phylogenies [46]. As expected, however, and similar to *Wolbachia*-infected *An. gambiae* s.l. from Burkina Faso [39] and Mali [40], a fragment of the wsp gene was not amplified from *Wolbachia-positive* samples from *An. gambiae* s.s, *An. arabiensis* and *An. coluzzii.* Similarly, no wsp gene fragment amplification occurred from *w*AnM-infected *An. moucheti.* However, wsp sequences were obtained from both *Wolbachia*-infected individuals of *An.* species A from Katana. We also analysed the wsp sequences of 22 specimens of *An.* species A from Lwiro (near Katana) and found identical sequences to the two individuals from Katana. Phylogenetic analysis of the wsp sequences obtained for the *w*AnsA strain, for both individuals from Katana (*w*AnsA wsp DRC-KAT1, *w*AnsA wsp DRC-KAT2) and three representative individuals from Lwiro (*w*AnsA wsp DRC-LWI1, *w*AnsA wsp DRC-LWI2, *w*AnsA wsp DRC-LWI3) indicates *w*AnsA is most closely related to *Wolbachia* strains of Supergroup B (such as *w*Pip, *w*AlbB, *w*Ma and *w*No) which is consistent with 16S rRNA phylogeny. However, the improved phylogenetic resolution provided by wsp indicates they cluster separately (**Figure 4b**). Typing of the *w*AnsA wsp nucleotide sequences highlighted that there were no exact matches to wsp alleles currently in the *Wolbachia* MLST database (https://pubmlst.org/wolbachia/) (**Table 2**). All *Wolbachia* 16S and wsp sequences were deposited into GenBank (accession numbers MH605275 – MH605285) **(Supplementary Table 2)**.

**Table 2.** Novel resident *Wolbachia* strain wsp and MLST gene allelic profiles. Exact matches to existing alleles present in the database are shown in bold, novel alleles are denoted by the allele number of the closest match and shown in red (number of single nucleotide differences to the closest match). *alternative degenerate primers (set 3) used to generate sequence.

| Mosquito species | *Wolbachia* strain | *Wolbachia* gene allele |  |  |  |  |  |
| --- | --- | --- | --- | --- | --- | --- | --- |
|  |  | *wsp* | *gatB* | *coxA* | *hcpA* | *ftsZ* | *fbpA* |
| *An. species A* | wAnsA | 152 (34) | 140 (4) | 122 (16) | 6 (7) | 7 (1) | 10 (1) |
| *An. species A* | wAnsA(2) | 152 (34) | - | 36 (1) | 6 (7) | - | - |
| *An. moucheti* | wAnM | - | 9 (2) | 11 (1) | 74 (3) | 7 (2) | 7 (12) |
| *An. coluzzii* | wAnga-Ghana | - | 9 | 64 | 3* | 177 | 4 |

**Figure 4.**
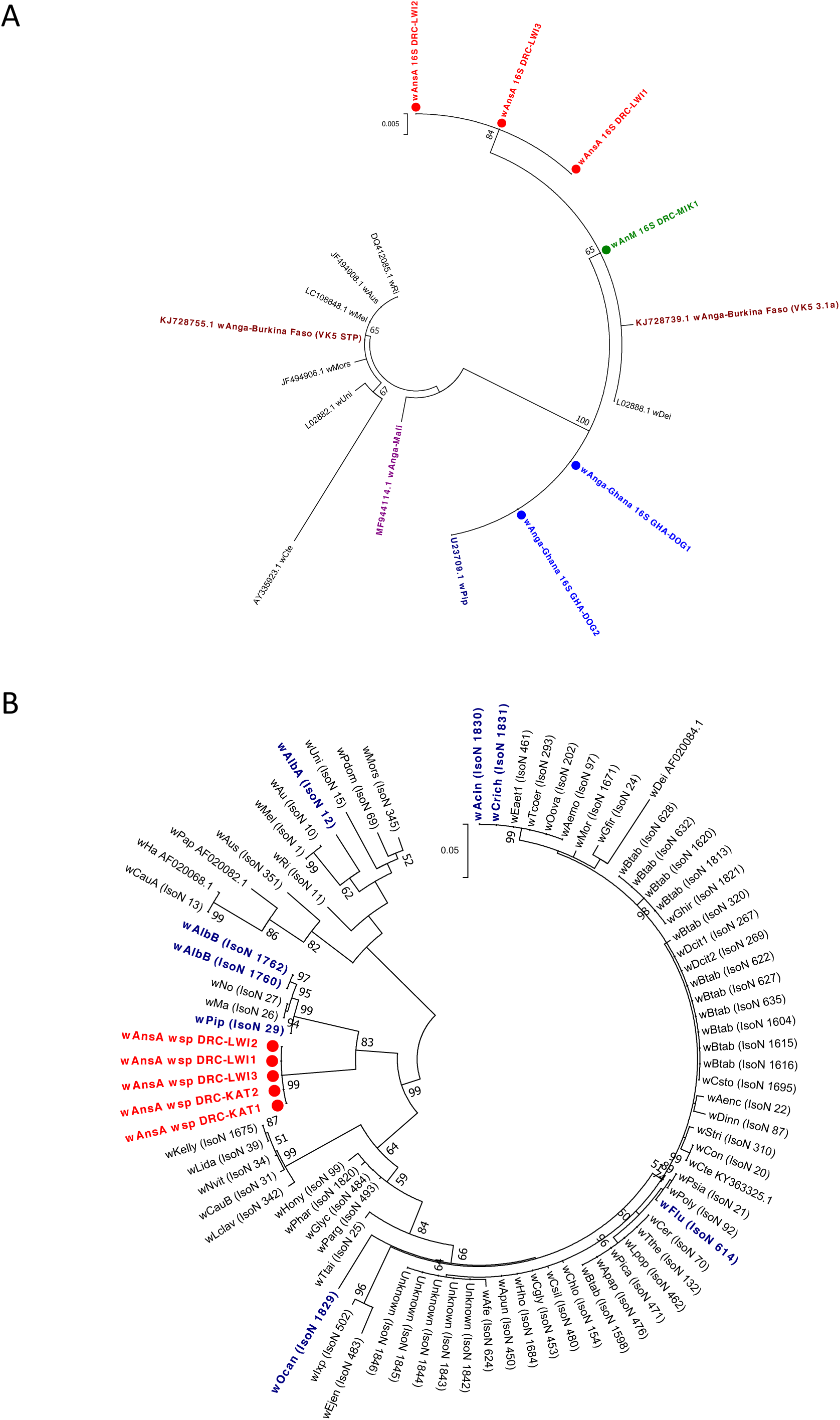
Resident *Wolbachia* strain phylogenetic analysis using 16S rRNA and wsp genes. **A)** Maximum Likelihood molecular phylogenetic analysis of the 16S rRNA gene for resident strains in *An. coluzzii* (*w*Anga-Ghana; blue), *An. moucheti* (*w*AnM; green) and *An. species A* (*w*AnsA; red). The tree with the highest log likelihood (-660.03) is shown. The tree is drawn to scale, with branch lengths measured in the number of substitutions per site. The analysis involved 17 nucleotide sequences. There were a total of 333 positions in the final dataset. Accession numbers of additional sequences obtained from GenBank are shown, including *w*Pip (navy blue), *w*Anga-Mali (purple) and *w*Anga-Burkina Faso strains (maroon). **B)** Maximum Likelihood molecular phylogenetic analysis of the wsp gene for *w*AnsA-infected representative individuals from the DRC (red). (KAT = Katana, LWI = Lwiro.) The tree with the highest log likelihood (-3663.41) is shown. The tree is drawn to scale, with branch lengths measured in the number of substitutions per site. The analysis involved 83 nucleotide sequences. There were a total of 443 positions in the final dataset. Reference numbers of additional sequences obtained from the MLST database (IsoN = Isolate number) or GenBank (accession number) are shown. Strains isolated from mosquitoes are highlighted in navy blue.

Multilocus sequence typing (MLST) was undertaken to provide more accurate strain phylogenies. This was done for the novel *Wolbachia* strains *w*AnM and *w*AnsA in addition to the resident *w*Anga-Ghana strain in *An. coluzzii* from Ghana. We were unable to amplify any of the five MLST genes from *Wolbachia*-infected *An. gambiae* s.s. and *An. arabiensis* from DRC (likely due to low infection densities). New alleles for all five MLST gene loci (sequences differed from those currently present in the MLST database) confirm the diversity of these novel *Wolbachia* strains (**Table 2**). The phylogeny of these three novel strains based on concatenated sequences of all five MLST gene loci confirms they cluster within Supergroup B (**Figure 5a**). This also demonstrates the novelty as comparison with a wide range of strains (including all isolates highlighted through partial matching during typing of each locus) shows these strains are distinct from currently available sequences (**Figure 5a, Table 2**). The concatenated phylogeny indicates that *w*AnM is most closely related to a Hemiptera strain: Isolate number 1616 found in *Bemisia tabaci* in Uganda, and a Coleoptera strain: Isolate number 20 found in *Tribolium confusum.* Concatenation of the MLST loci also indicates *w*AnsA is closest to a group containing various Lepidoptera and Hymenoptera strains from multiple countries in Asia, Europe and America, as well as two mosquito strains: Isolate numbers 1830 and 1831, found in *Aedes cinereus* and *Coquillettidia richiardii* in Russia. This highlights the lack of concordance between *Wolbachia* strain phylogeny and their insect hosts across diverse geographical regions. We also found evidence of potential strain variants in *w*AnsA through variable MLST gene fragment amplification and resulting closest-match allele numbers. A second *w*AnsA-infected sample, *An. sp.* A/1 (W+) DRC-KAT2, only amplified hcpA and coxA gene fragments and although identical sequences were obtained for wsp (**Figure 4b**) and hcpA, genetic diversity was seen in the coxA sequences, with typing revealing a different, but still novel allele for the coxA sequence from this individual (*w*AnsA(2) coxA DRC-KAT2) (**Figure 5b**). MLST gene fragment amplification was also variable for *w*Anga-Ghana-infected *An. coluzzii*, requiring two individuals to generate the five MLST gene sequences, and for the hcpA locus, more degenerate primers (hcpA_F3/hcpA_R3) were required to generate sequence of sufficient quality for analysis. This is likely due to the low density of this strain potentially influencing the ability to successfully amplify all MLST genes, in addition to the possibility of genetic variation in primer binding regions. Despite the sequences generated for this strain producing exact matches with alleles in the database for each of the five gene loci, the resultant allelic profile, and therefore strain type, did not produce a match, showing this *w*Anga-Ghana strain is also a novel strain type. The closest matches to the *w*Anga-Ghana allelic profile were with strains from two Lepidopteran species: Isolate number 609 found in *Fabriciana adippe* from Russia, and Isolate number 658 found in *Pammene fasciana* from Greece, but each of these only produced a match for 3 out of the 5 loci. The concatenated phylogeny for this strain (**Figure 5a**) indicates that across the 5 MLST loci, *w*Anga-Ghana is actually most closely related to a Lepidopteran strain found in *Thersamonia thersamon* in Russia (Isolate number 132). The phylogeny of *Wolbachia* strains based on the coxA gene (**Figure 5b**) highlights the genetic diversity of both the *w*AnsA strain variants and also *w*Anga-Ghana compared to the *w*Anga-Mali strain [40]; coxA gene sequences are not available for *w*Anga strains from Burkina Faso [39]. All *Wolbachia* MLST sequences were deposited into GenBank (accession numbers MH605286 – MH605305) **(Supplementary Table 3)**.

**Figure 5.**
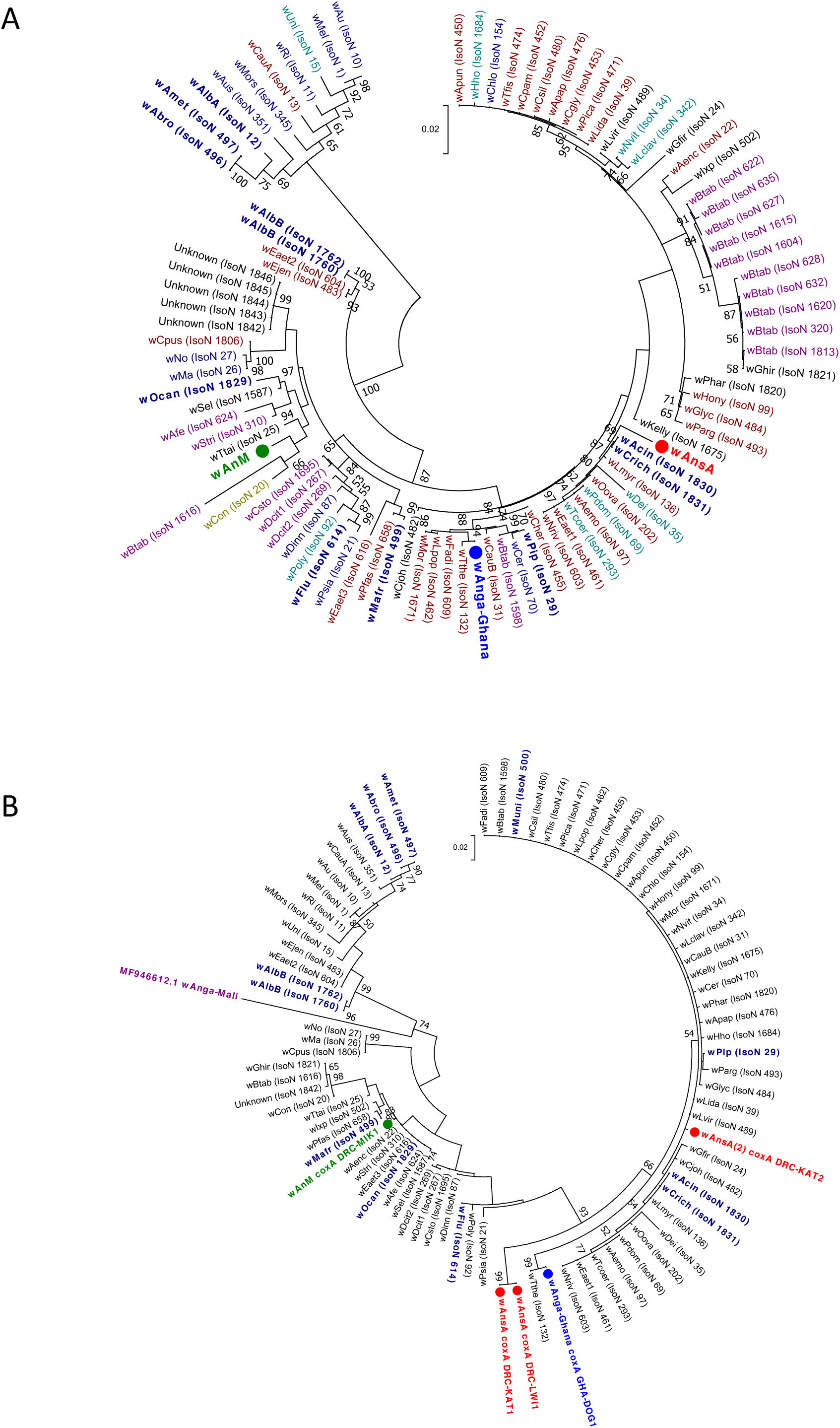
*Wolbachia* MLST phylogenetic analysis of resident *Wolbachia* strains in *An. coluzzii, An. moucheti* and *An.* species A. **A)** Maximum Likelihood molecular phylogenetic analysis from concatenation of all five MLST gene loci for resident *Wolbachia* strains from *An. coluzzii* (*w*Anga-Ghana; blue), *An. moucheti* (*w*AnM; green) and *An.* species A (*w*AnsA; red). The tree with the highest log likelihood (-10606.13) is shown and drawn to scale, with branch lengths measured in the number of substitutions per site. The analysis involved 94 nucleotide sequences. There were a total of 2067 positions in the final dataset. Concatenated sequence data from *Wolbachia* strains downloaded from MLST database for comparison shown with isolate numbers in brackets (IsoN). *Wolbachia* strains isolated from mosquito species highlighted in navy blue, bold. Strains isolated from other Dipteran species are shown in navy blue, from Coleoptera in olive green, from Hemiptera in purple, from Hymenoptera in teal blue, from Lepidoptera in maroon and from other, or unknown orders in black. **B)**. Maximum Likelihood molecular phylogenetic analysis for coxA gene locus for resident *Wolbachia* strains from *An. coluzzii* (*w*Anga-Ghana; blue), *An. moucheti* (*w*AnM; green) and *An.* species A (*w*AnsA and *w*AnsA(2); red). The tree with the highest log likelihood (-1921.11) is shown and drawn to scale, with branch lengths measured in the number of substitutions per site. The analysis involved 84 nucleotide sequences. There were a total of 402 positions in the final dataset. Sequence data for the coxA locus from *Wolbachia* strains downloaded from MLST database for comparison shown in black and navy blue with isolate numbers (IsoN) from MLST database shown in brackets. *Wolbachia* strains isolated from mosquito species highlighted in navy blue. GenBank sequence for *w*Anga-Mali coxA shown in maroon with accession number.

**Resident strain densities and relative abundance**. The relative densities of *Wolbachia* strains were estimated using qPCR targeting the ftsZ [47] and 16S rRNA [40] genes. ftsZ and 16S rRNA qPCR analysis indicated the amount of *Wolbachia* detected in *w*AnsA-infected and *w*AnM-infected females was approximately 1000-fold higher (Ct values 20-22) than *Wolbachia*-infected *An. gambiae s.s., An. arabiensis* and *w*Anga-Ghana-infected *An. coluzzii* (Ct values 30-33). To account for variation in mosquito body size and DNA extraction efficiency, we compared the total amount of DNA for *Wolbachia*-infected mosquito extracts and conversely, we found less total DNA in the *w*AnsA-infected extract (1.36 ng/μL) and the *An. moucheti* (*w*AnM-infected) extract (5.85 ng/μL) compared to the mean of 6.64 +/- 2.33 ng/μL for *w*Anga-Ghana-infected *An. coluzzii.* To estimate the relative abundance of resident *Wolbachia* strains in comparison to other bacterial species, we sequenced the bacterial microbiome using 16S rRNA amplicon sequencing on *Wolbachia*-infected individuals. We found *w*AnsA, *w*AnsA(2) and *w*AnM *Wolbachia* strains were the dominant operational taxonomic units (OTUs) of these mosquito species (**Figure 6**). In contrast, the lower density infection *w*Anga-Ghana strain represented only ~10% of the OTUs within the microbiome.

**Figure 6.**
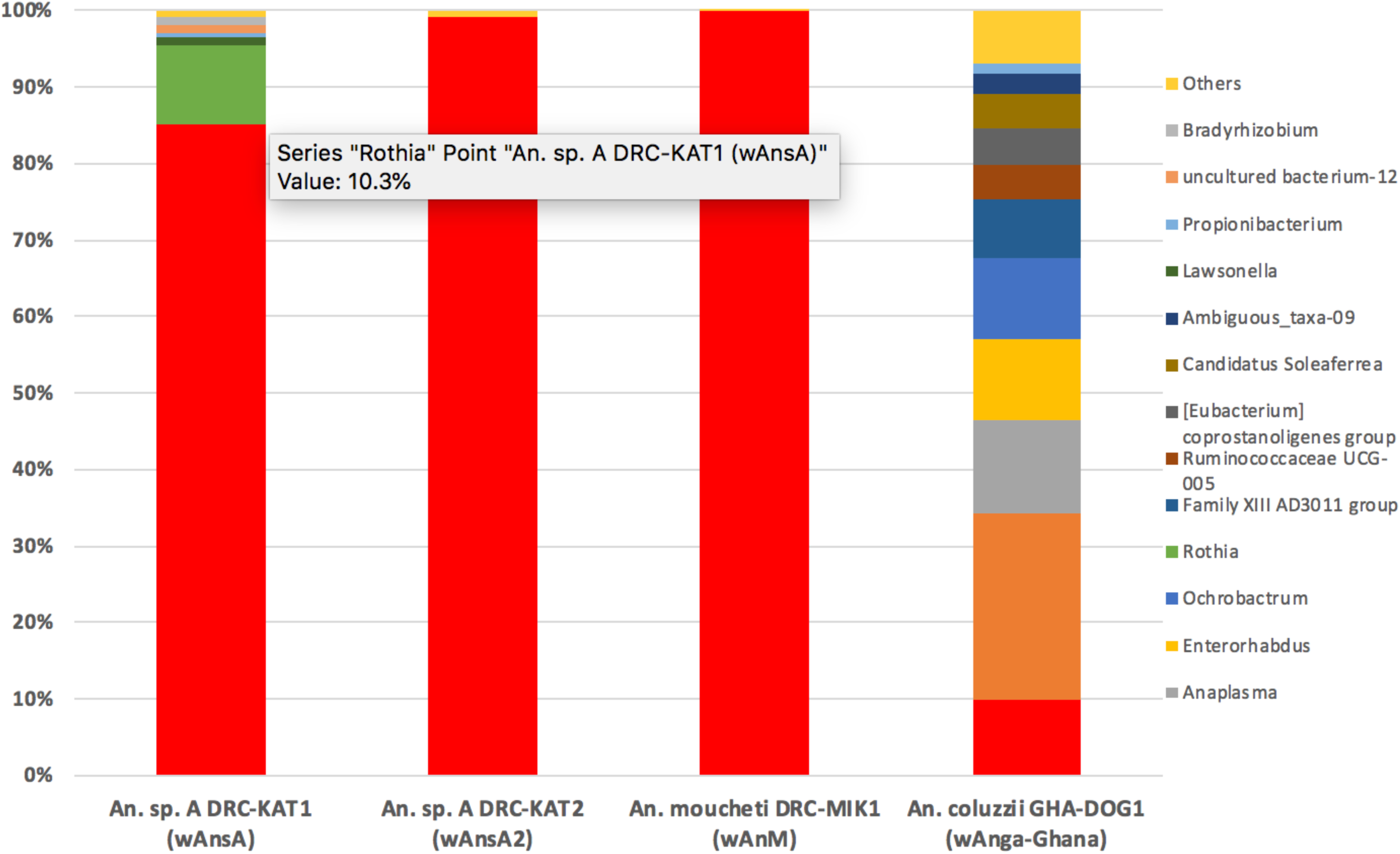
The relative abundance of resident *Wolbachia* strains in *Anopheles.* Bacterial genus level taxonomy was assigned to OTUs clustered with a 97% cut-off using the SILVA SSU v128 97% database, and individual genera comprising less than 1% of total abundance was merged into “Others”.

***P. falciparum, Wolbachia* and *Asaia* prevalence**. The prevalence of *P. falciparum* in female mosquitoes was extremely variable across countries and collection locations (**Figure 1, Table 1**) with very high prevalence recorded in *An. gambiae* s.s. from villages close to Boke (52%) and Faranah (44%) in Guinea. Despite the collection of other *Anopheles* species in Guinea, *An. gambiae* s.s. was the only species to have detectable malaria infections. In contrast, malaria was detected in multiple major vector species from DRC, including *An. gambiae* s.s, *An. arabiensis* and *An. funestus* s.s. A high prevalence of *P. falciparum* was also detected in *An. gambiae* s.s. from Uganda for both collection years; 19% for 2013 and 36% for 2014. In contrast, no *P. falciparum* infections were detected in any of the *An. coluzzii or An. melas* collected in Ghana. In Madagascar, *P. falciparum* was detected in only two species; *An. gambiae s.s.* and *An. rufipes.* We compared the overall *P. falciparum* infection rates in *An. gambiae* s.s. mosquitoes collected across all locations from DRC to determine if there was any correlation with the presence of the low density *w*Anga-DRC *Wolbachia* resident strain. Overall, of the 128 mosquitoes collected, only 1.56% (n=2) had detectable *Wolbachia-Plasmodium* co-infections compared to 10.16% (n=13) where we only detected *Wolbachia.* A further 11.72% (n=15) were only PCR-positive for *P. falciparum.* As expected, for the vast majority of mosquitoes (76.56%, n=98) we found no evidence of *Wolbachia* or *P. falciparum* present, resulting in no correlation across all samples (Fisher’s exact *post hoc* test on unnormalized data, two-tailed, *P*=0.999). Interestingly, one *An.* species ‘A’ female from Katana was infected with *P. falciparum.*

For all *Wolbachia*-infected females collected in our study (including *An. coluzzii* from Ghana and novel resident strains in *An. moucheti* and *An. species* A), we did not detect the presence of *Asaia.* No resident *Wolbachia* strain infections were detected in *Anopheles* mosquitoes from Guinea, Uganda or Madagascar. However, high *Asaia* and malaria prevalence rates were present in *Anopheles* mosquitoes from Uganda and Guinea (including multiple species in all four sites in Guinea). We compared the overall *P. falciparum* infection rates in *An. gambiae* s.s. collected across all locations from Guinea, with and without *Asaia* bacteria, and found no overall correlation (Fisher’s exact *post hoc* test on unnormalized data, two-tailed, *P*=0.4902). There was also no overall correlation between *Asaia* and *P. falciparum* infections in *An. gambiae* s.s. from Uganda for both 2013 (Fisher’s exact *post hoc* test on unnormalized data, two-tailed, *P*=0.601) and 2014 (Fisher’s exact *post hoc* test on unnormalized data, two-tailed, *P*=0.282).

*Asaia* can be environmentally acquired at all life stages but can also have the potential to be vertically and horizontally transmitted between individual mosquitoes. Therefore, we performed 16S microbiome analysis on a sub-sample of *Asaia*-infected *An. gambiae* s.s. from Kissidougou (Guinea), a location in which high levels of *Asaia* were detected by qPCR (mean *Asaia* Ct = 17.84 +/- 2.27). *Asaia* in these individuals is the dominant bacterial species present (**Figure 7a**) but in Uganda we detected much lower levels of *Asaia* (qPCR mean Ct = 33.33 +/- 0.19) and this was reflected in *Asaia* not being a dominant species (**Figure 7b**). The alpha and beta diversity of *An. gambiae* s.s. from Kissidougou, Guinea and Butemba, Uganda shows much more overall diversity in the microbiome for Uganda individuals (**supplementary figure S1**). Interestingly, 2/5 of these individuals from Kissidougou (Guinea) were *P. falciparum-infected* compared to 3/5 individuals from Uganda. To determine if the presence of *Asaia* had a quantifiable effect on the level of *P. falciparum* detected, we normalized *P. falciparum* Ct values from qPCR (**supplementary figure S2a**) and compared gene ratios for *An. gambiae s.s.* mosquitoes from Guinea, with or without *Asaia* (**supplementary figure S2b**). Statistical analysis using student’s t-tests revealed no significant difference between normalized *P. falciparum* gene ratios (p= 0.51, df =59). Larger variation of Ct values was seen for *Asaia* (**supplementary figure S2c**) suggesting the bacterial densities in individual mosquitoes were more variable than *P. falciparum* parasite infection levels.

**Figure 7.**
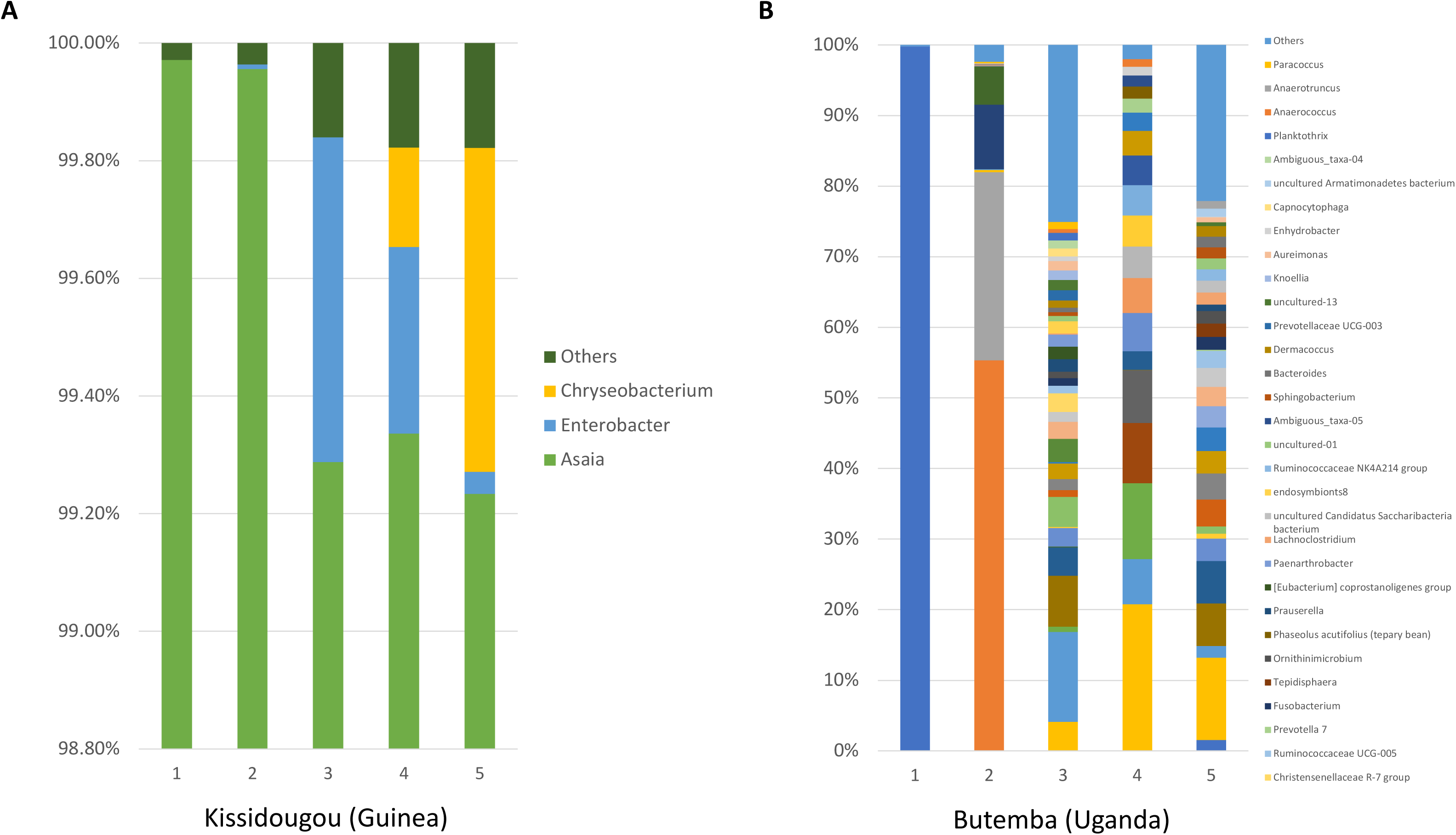
The relative abundance of bacteria in *An. gambiae* s.s. comparing two locations with contrasting *Asaia* infection densities. Bacterial genus level taxonomy was assigned to OTUs clustered with a 97% cut-off using the SILVA SSU v128 97% database, and individual genera comprising less than 1% of total abundance was merged into “Others”.

### Discussion

Malaria transmission in Sub-Saharan Africa is highly dependent on the local *Anopheles* vector species but the primary vector complexes recognised are *An. gambiae* s.l., *An. funestus* s.l. *An. nili* s.l. and *An. moucheti* s.l. [41,48]. *An. gambiae* s.s. and *An. coluzzii* sibling species are considered the most important malaria vectors in Sub-Saharan Africa and recent studies indicate that *An. coluzzii* extends further north, and closer to the coast than *An. gambiae s.s.* within west Africa [49]. In our study, high malaria prevalence rates in *An. gambiae* s.s. across Guinea would be consistent with high malaria parasite prevalence (measured by rapid diagnostic tests) in Guéckédou prefecture, and the overall national malaria prevalence estimated to be 44% in 2013 [50]. However, malaria prevalence has decreased in the past few years with an overall prevalence across Guinea estimated at 15% for 2016. Although our *P. falciparum* infection prevalence rates were also high in DRC, recent studies have shown comparable levels of infection with 35% of *An. gambiae* s.l. mosquitoes infected from Kinshasa [51]. We detected *P. falciparum* in *An. gambiae* s.s, *An. arabiensis, An. funestus* s.s. and *An.* species A from DRC. Morphological differences have been widely used for identification of malaria vectors but species complexes (such as *An. gambiae* s.l. and *An. funestus* s.l.) require species-diagnostic PCR assays. Historically, malaria entomology studies in Africa have focused predominantly on species from these complexes, likely due to the fact that mosquitoes from these complexes dominate the collections [43]. In our study, we used ITS2 sequencing to confirm secondary vector species that were *P. falciparum-infected* given the difficulties of morphological identification and recent studies demonstrating the inaccuracy of diagnostic species PCR-based molecular identification [52]. Our study is the first to report the detection of *P. falciparum* in *An. rufipes* from Madagascar; previously this species was considered a vector of *Plasmodium* species of non-human origin and has only very recently been implicated in human malaria transmission [53]. However, detection of *P. falciparum* parasites in whole body mosquitoes does not confirm that the species plays a significant role in transmission. Detection could represent infected bloodmeal stages or oocysts present in the midgut wall so further studies are warranted to determine this species ability to transmit human malaria parasites.

The mosquito microbiota can modulate the mosquito immune response and bacteria present in wild *Anopheles* populations can influence malaria vector competence [4,5]. Endosymbiotic *Wolbachia* bacteria are particularly widespread through insect populations but they were commonly thought to be absent from *Anopheles* mosquitoes. However, the recent discovery of *Wolbachia* strains in the *An. gambiae* s.l. complex in Burkina Faso and Mali [39,40] in addition to our study showing infection in *Anopheles* from Ghana and DRC, suggest resident strains could be widespread across Sub-Saharan Africa. The discovery of resident strains in Burkina Faso resulted from sequencing of the 16S rRNA gene identifying *Wolbachia* sequences rather than screening using *Wolbachia-specific* genes [39]. Intriguingly, *Wolbachia* infections in these mosquitoes could not be detected using conventional PCR targeting the wsp gene. As the wsp gene has often been used in previous studies to detect strains in *Anopheles* species [25,27], this could explain why resident strains in the *An. gambiae* s.l. complex have gone undetected until very recently. Recent similar methods using 16S rRNA amplicon sequencing to determine the overall microbiota in wild mosquito populations has provided evidence for *Wolbachia* infections in *An. gambiae* in additional villages in Burkina Faso [54] and *Anopheles* species collected in Illinois, USA [55]. Our study describing resident *Wolbachia* strains in numerous species of *Anopheles* malaria vectors also highlights the potential for *Wolbachia* to be influencing malaria transmission, as postulated by previous studies [39,40,56]. Although no significant correlation was present for malaria and *Wolbachia* prevalence in the 128 *An. gambiae* s.s. individuals from DRC, we only detected co-infections in two individuals compared to 13 and 15 individuals infected only with *Wolbachia* or *P. falciparum* respectively. As the majority (77%) of samples had neither detectable *Wolbachia* resident strains or *P. falciparum*, a larger sample size would be needed to determine if there is a correlation, as shown previously in both Burkina Faso [56] and Mali [40]. The infection prevalence of resident *Wolbachia* strains in *An. coluzzii* from Ghana (4%) and *An. gambiae* s.s. from the DRC was variable but low (8-24%), consistent with infection prevalence in Burkina Faso (11%) [39] but much lower than those reported in Mali (60-80%) [40] where infection was associated with reduced prevalence and intensity of sporozoite infection in field-collected females.

The discovery of a resident *Wolbachia* strain in *An. moucheti*, a highly anthropophilic and efficient malaria vector found in the forested areas of western and central Africa [41], suggests further studies are warranted that utilize large sample sizes to examine the influence of the *w*AnM *Wolbachia* strain on *Plasmodium* infection dynamics in this malaria vector. *An. moucheti* is often the most abundant vector, breeding in slow moving streams and rivers, contributing to year round malaria transmission in these regions [57,58]. This species has also been implicated as a main bridge vector species in the transmission of ape *Plasmodium* malaria in Gabon [59]. There is thought to be high genetic diversity in *An. moucheti* populations [60,61] which may either influence the prevalence of *Wolbachia* resident strains or *Wolbachia* could be contributing to genetic diversity through its effect on host reproduction. A novel *Wolbachia* strain in *An.* species ‘A’, present at high infection frequencies in Lwiro (close to Katana in DRC), also suggests more *Anopheles* species, including unidentified and potentially new species, could be infected with this widespread endosymbiotic bacterium. *An.* species A should be further investigated to determine if this species is a potential malaria vector given our study demonstrated *P. falciparum* infection in one of two individuals screened and ELISA-positive samples of this species were reported from the Western Highlands of Kenya [62].

The variability of *Wolbachia* prevalence rates in *An. gambiae* s.l. complex from locations within DRC and Ghana and previous studies in Burkina Faso [39] and Mali [40] suggest the environment is one factor that influences the presence or absence of resident strains. In our study we found no evidence of *Wolbachia-Asaia* co-infections across all countries, supporting laboratory studies that have shown these two bacterial endosymbionts demonstrate competitive exclusion in *Anopheles* species [36,38]. We also found that *Asaia* infection densities (whole body mosquitoes) were variable and location dependent which would correlate with this bacterium being environmentally acquired at all life stages, but also having the potential for both vertical and horizontal transmission [37]. Significant variations in overall *Asaia* prevalence and density across different *Anopheles* species and locations in our study would also correlate with our data indicating no evidence of an association with *P. falciparum* prevalence in both Guinea and Uganda populations. Further studies are needed to determine the complex interaction between these two bacterial endosymbionts and malaria in diverse *Anopheles* malaria vector species. Horizontal transfer of *Wolbachia* strains between species (even over large phylogenetic differences) has shaped the evolutionary history of this endosymbiont in insects and there is evidence for loss of infection in host lineages over evolutionary time [63]. Our results showing a new strain present in *An. coluzzii* from Ghana (phylogenetically different to strains present in *An. gambiae* s.l. mosquitoes from both Burkina Faso and Mali), strain variants observed in An. species A, and the concatenated grouping of the novel *Anopheles* strains with strains found in different Orders of insects, support the lack of congruence between insect host and *Wolbachia* phylogenetic trees [64].

Our qPCR and 16S microbiome analysis indicates the densities of *w*AnM and *w*AnsA strains are significantly higher than resident *Wolbachia* strains in *An. gambiae* s.l. However, caution must be taken as we were only able to analyse selected individuals and larger collections of wild populations would be required to confirm these results. Native *Wolbachia* strains dominating the microbiome of *An. species* A and *An. moucheti* is consistent with other studies of resident strains in mosquitoes showing *Wolbachia* 16S rRNA gene amplicons vastly outnumber sequences from other bacteria in *Ae. albopictus* and *Cx. quinquefasciatus* [65,66]. The discovery of novel *Wolbachia* strains provides the rationale to undertake vector competence experiments to determine what effect these strains are having on malaria transmission. The tissue tropism of novel *Wolbachia* strains in malaria vectors will be particularly important to characterise given this will determine if these endosymbiotic bacteria are proximal to malaria parasites within the mosquito. It would also be important to determine the additional phenotypic effects novel resident *Wolbachia* strains have on their mosquito hosts. Some *Wolbachia* strains induce a reproductive phenotype termed cytoplasmic incompatibility (CI) that results in inviable offspring when an uninfected female mates with a *Wolbachia*-infected male. In contrast, *Wolbachia*-infected females produce viable progeny when they mate with both infected and uninfected male mosquitoes. This reproductive advantage over uninfected females allows *Wolbachia* to spread within mosquito populations.

*Wolbachia* has been the focus of recent biocontrol strategies in which *Wolbachia* strains transferred into naïve mosquito species provide strong inhibitory effects on arboviruses [19,20,67–70] and malaria parasites [31,35]. The discovery of two novel *Wolbachia* strains in *Anopheles* mosquitoes, potentially present at much higher density than resident strains in the *An. gambiae* s.l. complex, also suggests the potential for these strains to be transinfected into other *Anopheles* species to produce inhibitory effects on *Plasmodium* parasites. *Wolbachia* transinfection success is partly attributed to the relatedness of donor and recipient host so the transfer of high density *Wolbachia* strains between *Anopheles* species may result in stable infections (or co-infections) that have strong inhibitory effects on *Plasmodium* development. Finally, if the resident strain present in *An. moucheti* is at low infection frequencies in wild populations, an alternative strategy known as the incompatible insect technique (IIT) could be implemented where *Wolbachia*-infected males are released to suppress the wild populations through CI (reviewed by [22]). In summary, the important discovery of diverse novel *Wolbachia* strains in *Anopheles* species will help our understanding of how *Wolbachia* strains can potentially impact malaria transmission, through natural associations or being used as candidate strains for transinfection to create stable infections in other species.

## Materials and Methods

### Study sites & collection methods

*Anopheles* adult mosquitoes were collected from five malaria endemic countries in Sub-Saharan Africa; Guinea, Democratic Republic of the Congo (DRC), Ghana, Uganda and Madagascar between 2013 and 2017 (**Figure 1**). Human landing catches, CDC light traps and pyrethrum spray catches were undertaken between April 2014 – February 2015 in 10 villages near four cities in Guinea; Foulayah (10.144633, -10.749717) and Balayani (10.1325, - 10.7443) near Faranah; Djoumaya (10.836317, -14.2481) and Kaboye Amaraya (10.93435, - 14.36995) near Boke; Tongbekoro (9.294295, -10.147953), Keredou (9.208919, -10.069525), and Gbangbadou (9.274363, -9.998639) near Kissidougou; and Makonon (10.291124, -9.363358), Balandou (10.407669, -9.219096), and Dalabani (10.463692, -9.451904) near Kankan. Human landing catches and pyrethrum spray catches were undertaken between January – September 2015 in seven sites of the DRC; Kinshasa (-4.415881, 15.412188), Mikalayi (-6.024184, 22.318251), Kisangani (0.516350, 25.221176), Katana (-2.225129, 28.831604), Kalemie (-5.919054, 29.186572), and Kapolowe (-10.939802, 26.952970). We also analysed a subset from collections obtained from Lwiro (-2.244097, 28.815232), a village near Katana, collected between in September – October 2015. A combination of CDC light traps, pyrethrum spray catches and human landing catches were undertaken in Butemba, Kyankwanzi District in mid-western Uganda (1.1068444, 31.5910085) in August and September 2013 and June 2014. CDC light trap catches were undertaken in May 2017 in Dogo in Ada, Greater Accra, Ghana (5.874861111, 0.560611111). In Madagascar, sampling was undertaken in June 2016 at four sites: Anivorano Nord, located in the Northern domain, (-12.7645000, 49.2386944), Ambomiharina, Western domain, (-16.3672778, 46.9928889), Antafia, Western domain, (-17.0271667, 46.7671389) and Ambohimarina, Central domain, (-18.3329444, 47.1092500). Trapping consisted of CDC light traps and a net trap baited with Zebu (local species of cattle) to attract zoophilic species [71].

### DNA extraction and species identification

DNA was extracted from individual whole mosquitoes or abdomens using QIAGEN DNeasy Blood and Tissue Kits according to manufacturer’s instructions. DNA extracts were eluted in a final volume of 100 μL and stored at −20°C. Species identification was initially undertaken using morphological keys followed by diagnostic species-specific PCR assays to distinguish between the morphologically indistinguishable sibling mosquito species of the *An. gambiae* [72–74] and *An. funestus* complexes [75]. To determine species identification for samples of interest and samples that could not be identified by species-specific PCR, Sanger sequences were generated from ITS2 PCR products [76].

### Detection of P. falciparum and Asaia

Detection of *P. falciparum* malaria was undertaken using qPCR targeting an 120-bp sequence of the *P. falciparum* cytochrome c oxidase subunit 1 (Cox1) mitochondrial gene [77] as preliminary trials revealed this was the optimal method for both sensitivity and specificity. Positive controls from gDNA extracted from a cultured *P. falciparum-infected* blood sample (parasitaemia of ~10%) were serially diluted to determine the threshold limit of detection, in addition to the inclusion no template controls (NTCs). *Asaia* detection was undertaken targeting the 16S rRNA gene [78,79]. Ct values for both *P. falciparum* and *Asaia* assays in selected *An. gambiae* extracts were normalized to Ct values for a single copy *An. gambiae* rps17 housekeeping gene (accession no. AGAP004887 on www.vectorbase.org) [80,81]. As Ct values are inversely related to the amount of amplified DNA, a higher target gene Ct: host gene Ct ratio represented a lower estimated infection level. qPCR reactions were prepared using 5 μL of FastStart SYBR Green Master mix (Roche Diagnostics), a final concentration of 1μM of each primer, 1 μL of PCR grade water and 2 μL template DNA, to a final reaction volume of 10 μL. Prepared reactions were run on a Roche LightCycler^®^ 96 System and amplification was followed by a dissociation curve (95°C for 10 seconds, 65°C for 60 seconds and 97°C for 1 second) to ensure the correct target sequence was being amplified. PCR results were analysed using the LightCycler^®^ 96 software (Roche Diagnostics). A sub-selection of PCR products from each assay was sequenced to confirm correct amplification of the target gene fragment.

### Wolbachia detection

*Wolbachia* detection was first undertaken targeting three conserved *Wolbachia* genes previously shown to amplify a wide diversity of strains; 16S rDNA gene [40,45], *Wolbachia* surface protein (wsp) gene [46] and FtsZ cell cycle gene [82]. DNA extracted from a *Drosophila melanogaster* fly (infected with the *w*Mel strain of *Wolbachia*) was used a positive control, in addition to no template controls (NTCs). 16S rDNA [45] and wsp [46] gene PCR reactions were carried out in a Bio-Rad T100 Thermal Cycler using standard cycling conditions and PCR products were separated and visualised using 2% E-Gel EX agarose gels (Invitrogen) with SYBR safe and an Invitrogen E-Gel iBase Real-Time Transilluminator. FtsZ [47] and *16S* rDNA [40] gene real time PCR reactions were prepared using 5 μL of FastStart SYBR Green Master mix (Roche Diagnostics), a final concentration of 1μM of each primer, 1 μL of PCR grade water and 2 μL template DNA, to a final reaction volume of 10 μL. Prepared reactions were run on a Roche LightCycler^®^ 96 System for 15 minutes at 95°C, followed by 40 cycles of 95°C for 15 seconds and 58°C for 30 seconds. Amplification was followed by a dissociation curve (95°C for 10 seconds, 65°C for 60 seconds and 97°C for 1 second) to ensure the correct target sequence was being amplified. PCR results were analysed using the LightCycler^®^ 96 software (Roche Diagnostics). To estimate *Wolbachia* densities across multiple *Anopheles* mosquito species, *ftsZ* and 16S qPCR Ct values were compared to total dsDNA extracted measured using an Invitrogen Qubit 4 fluorometer. A serial dilution series of a known *Wolbachia*-infected mosquito DNA extract was used to correlate Ct values and amount of amplified target product.

### Wolbachia MLST

Multilocus sequence typing (MLST) was undertaken to characterize *Wolbachia* strains using the sequences of five conserved genes as molecular markers to genotype each strain. In brief, 450-500 base pair fragments of the gatB, coxA, hcpA, ftsZ and fbpA *Wolbachia* genes were amplified from individual *Wolbachia*-infected mosquitoes using previously optimised protocols [83]. A *Cx. pipiens* gDNA extraction (previously shown to be infected with the *w*Pip strain of *Wolbachia*) was used a positive control for each PCR run, in addition to no template controls (NTCs). If no amplification was detected using standard primers, further PCR analysis was undertaken using degenerate primers [83]. PCR products were separated and visualised using 2% E-Gel EX agarose gels (Invitrogen) with SYBR safe and an Invitrogen E-Gel iBase Real-Time Transilluminator. PCR products were submitted to Source BioScience (Source BioScience Plc, Nottingham, UK) for PCR reaction clean-up, followed by Sanger sequencing to generate both forward and reverse reads. Sequencing analysis was carried out in MEGA7 [84] as follows. Both chromatograms (forward and reverse traces) from each sample was manually checked, edited, and trimmed as required, followed by alignment by ClustalW and checking to produce consensus sequences. Consensus sequences were used to perform nucleotide BLAST (NCBI) database queries, and searches against the *Wolbachia* MLST database (http://pubmlst.org/wolbachia) [85]. If a sequence produced an exact match in the MLST database we assigned the appropriate allele number, otherwise the closest matches and number of differences were noted. The Sanger sequencing traces from the wsp gene were also treated in the same way and analysed alongside the MLST gene locus scheme, as an additional marker for strain typing

### Phylogenetic analysis

Alignments were constructed in MEGA7 by ClustalW to include all relevant and available sequences highlighted through searches on the BLAST and *Wolbachia* MLST databases. Maximum Likelihood phylogenetic trees were constructed from Sanger sequences as follows. The evolutionary history was inferred by using the Maximum Likelihood method based on the Tamura-Nei model [86]. The tree with the highest log likelihood in each case is shown. The percentage of trees in which the associated taxa clustered together is shown next to the branches. Initial tree(s) for the heuristic search were obtained automatically by applying Neighbor-Join and BioNJ algorithms to a matrix of pairwise distances estimated using the Maximum Composite Likelihood (MCL) approach, and then selecting the topology with superior log likelihood value. The trees are drawn to scale, with branch lengths measured in the number of substitutions per site. Codon positions included were 1st+2nd+3rd+Noncoding. All positions containing gaps and missing data were eliminated. The phylogeny test was by Bootstrap method with 1000 replications. Evolutionary analyses were conducted in MEGA7 [84].

### Microbiome Analysis

The microbiomes of selected individual *Anopheles* were analysed using barcoded high-throughput amplicon sequencing of the bacterial 16S rRNA gene. Sequencing libraries for each isolate were generated using universal 16S rRNA V3-V4 region primers [87] in accordance with Illumina 16S rRNA metagenomic sequencing library protocols. The samples were barcoded for multiplexing using Nextera XT Index Kit v2. Sequencing was performed on an Illumina MiSeq instrument using a MiSeq Reagent Kit v2 (500-cycles). Quality control and taxonomical assignment of the resultant reads were performed using CLC Genomics Workbench 8.0.1 Microbial Genomics Module (http://www.clcbio.com). Low quality reads containing nucleotides with quality threshold below 0.05 (using the modified Richard Mott algorithm), as well as reads with two or more unknown nucleotides were removed from analysis. Additionally reads were trimmed to remove sequenced Nextera adapters. Reference based OTU picking was performed using the SILVA SSU v128 97% database [88]. Sequences present in more than one copy but not clustered to the database were then placed into de novo OTUs (97% similarity) and aligned against the reference database with 80% similarity threshold to assign the “closest” taxonomical name where possible. Chimeras were removed from the dataset if the absolute crossover cost was 3 using a k-mer size of 6. Alpha diversity was measured using Shannon entropy (OTU level).

### Statistical analysis

Fisher’s exact *post hoc* test in Graphpad Prism 7 was used to compare infection rates. Normalised qPCR Ct ratios were compared using unpaired t-tests in GraphPad Prism 7.

## Authors’ contributions

MK, JO, JB, EH, MLT, FNR, KK, DC, YB, FW, EZM, YAA, ARM, TAA performed field collections. CLJ, GGL, MK, JO, KS, EH & TW performed sample analysis. CLJ performed sequence analysis. GG, SH, KK, MP, YF and GLH performed 16S microbiome sample analysis. SRI, GLH and TW provided overall supervision. CLJ and TW wrote the initial draft.

## Acknowledgements

We would like to thank all the mosquito collectors and residents of the villages where collections took place. We would also like to thank John Gimnig, Bill Hawley and Barb Marston for reviewing our manuscript. CLJ and TW were supported by a Wellcome Trust /Royal Society grant awarded to TW (101285/Z/13/Z): http://www.wellcome.ac.uk; https://royalsociety.org. GLH is supported by NIH grants (R21AI124452 and R21AI129507), a University of Texas Rising Star award, the John S. Dunn Foundation Collaborative Research Award, the Robert J. Kleberg, Jr. and Helen C. Kleberg Foundation, and the Centers for Disease Control and Prevention (CDC) (Cooperative Agreement Number U01CK000512). The papers contents are solely the responsibility of the authors and do not necessarily represent the official views of the CDC or the Department of Health and Human Services. This work was also supported by a James W. McLaughlin postdoctoral fellowship at the University of Texas Medical Branch to SH. Field work in Uganda was funded by UK aid (through the Programme Partnership Arrangement grant to Malaria Consortium). YAA and ARM were supported by a NIH grant R01AI123074. SRI was funded by the U.S. President’s Malaria Initiative. The funders had no role in study design, data collection and analysis, decision to publish, or preparation of the manuscript.

---

**Supplementary Figure S1.**
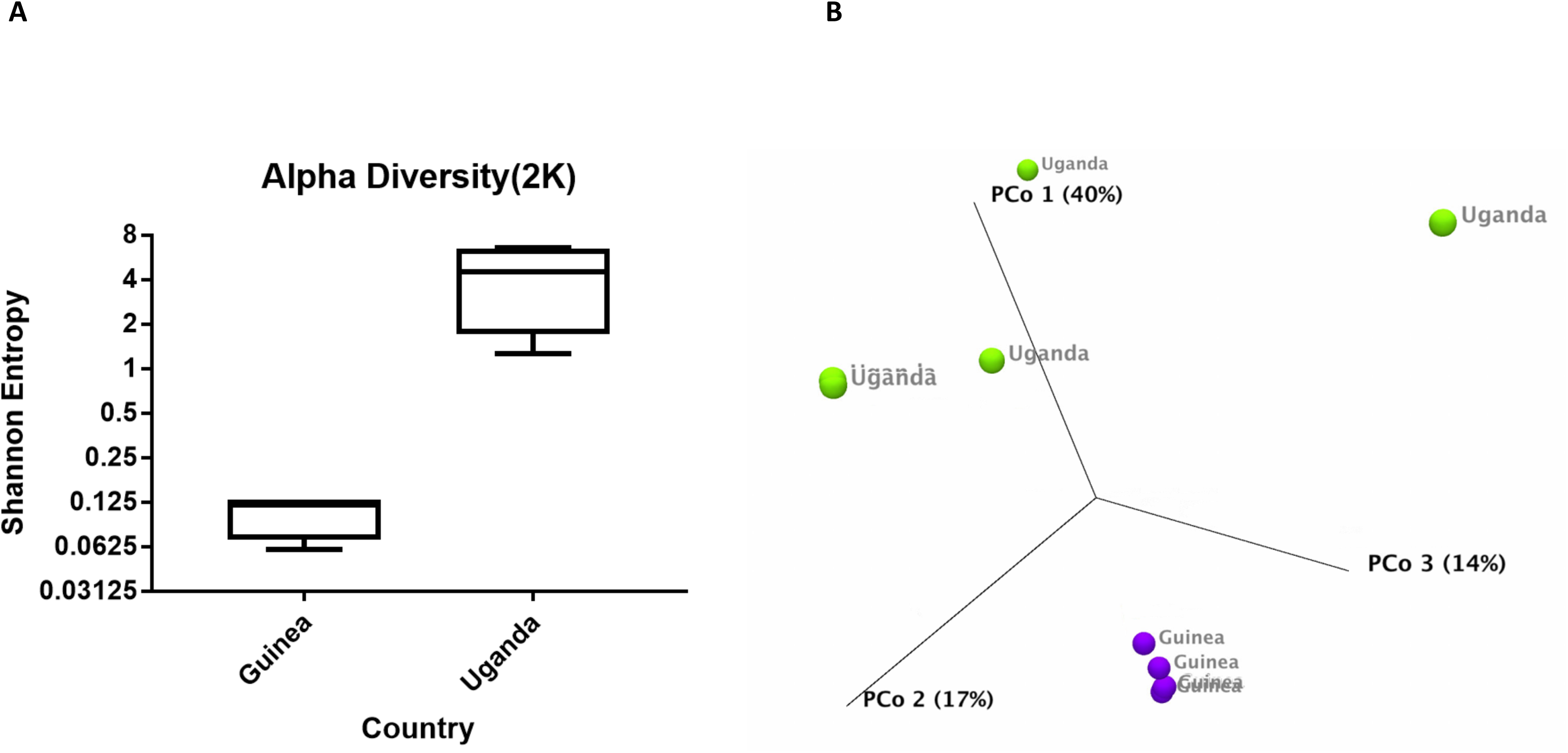
Alpha and beta diversity of *An. gambiae* s.s. from Kissidougou, Guinea and Butemba, Uganda. **A)** Alpha diversity using the Shannon diversity index shows the relative abundance of bacterial genera. **B)** To identify dissimilarities in the bacterial community structure between the microbiome, principal coordinates analysis (PCoA) was performed based on a Bray-Curtis dissimilarity matrix based on 97% clustered OTUs.

**Supplementary Figure S2.**
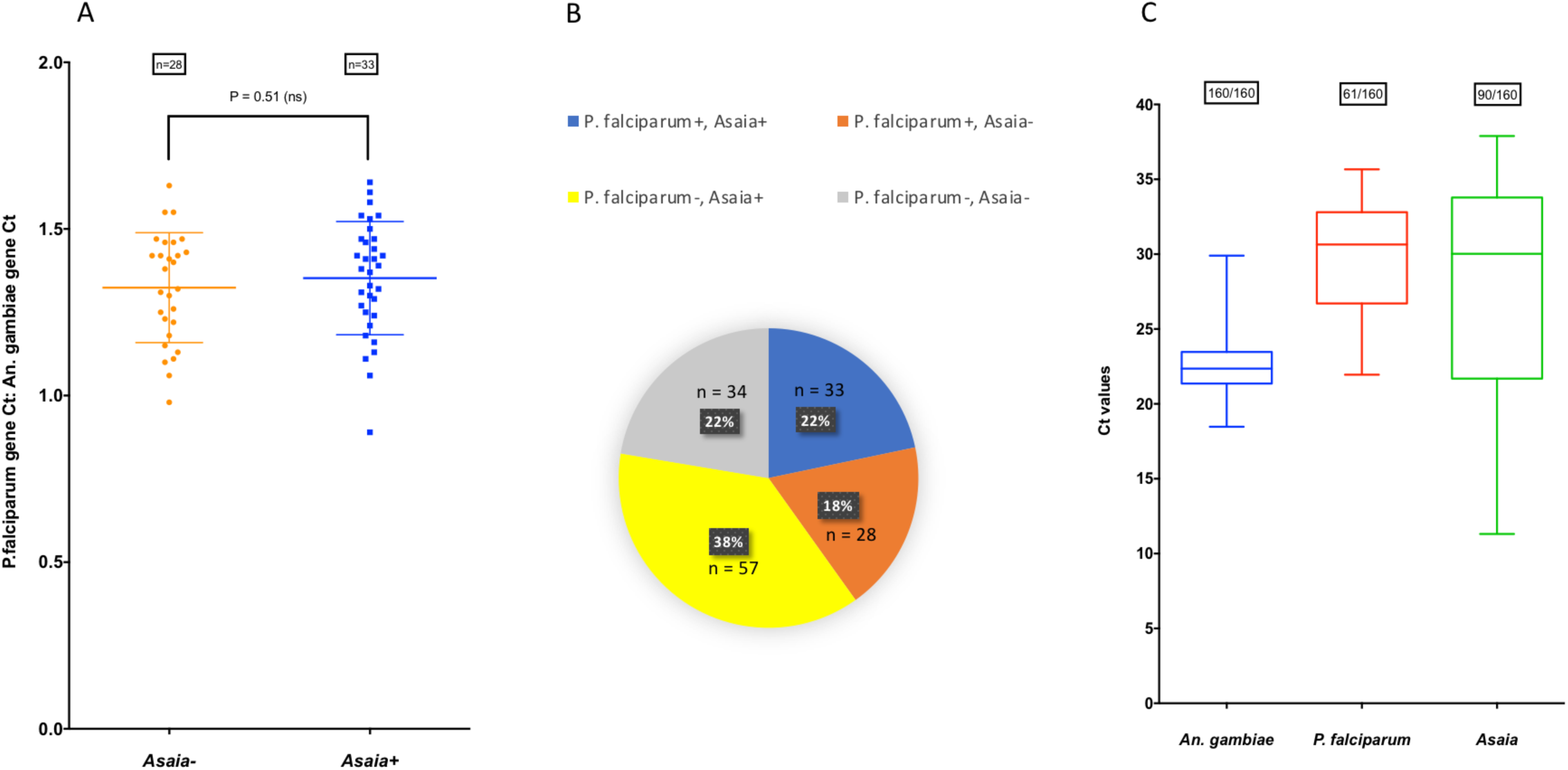
Prevalence of the bacterial endosymbiont *Asaia* and malaria parasites in *An. gambiae* s.s. mosquitoes from Guinea. **A)** Normalised *P. falciparum: An. gambiae* gene Ct ratio for mosquitoes that are infected with malaria and +/- *Asaia* bacteria. **B)** *P. falciparum* and *Asaia* infection rates (%) in 152 *An. gambiae* s.s. females. **C)** Box and whisker plot of Ct values for detection of *Asaia* and *P. falciparum* malaria showing more variable levels of *Asaia* detected.

**Supplementary Table 1.** Additional sample details and ITS2 GenBank accession numbers.

| Location | Species | Sample ID | *Wolbachia* status | ITS2 accession number |
| --- | --- | --- | --- | --- |
| Guinea: Kissidougou | *Anopheles* sp. O/15 | GUI-KSK1 | W- | MH598414 |
| Guinea: Kissidougou | *Anopheles gambiae* s.s. | GUI-KSK2 | W- | MH598415 |
| Guinea: Faranah | *Anopheles nili* | GUI-FAR1 | W- | MH598416 |
| Guinea: Faranah | *Anopheles nili* | GUI-FAR2 | W- | MH598417 |
| Guinea: Kankan | *Anopheles* sp. unknown | GUI-KAN1 | W- | MH598418 |
| DRC: Lwiro | *Anopheles* sp. A/1 | DRC-LWI1 | W+ | MH598419 |
| DRC: Lwiro | *Anopheles* sp. A/1 | DRC-LWI2 | W+ | MH598420 |
| DRC: Lwiro | *Anopheles* sp. A/1 | DRC-LWI3 | W+ | MH598421 |
| DRC: Katana | *Anopheles* sp. A/1 | DRC-KAT1 | W+ | MH598422 |
| DRC: Katana | *Anopheles* sp. A/1 | DRC-KAT2 | W+ | MH598423 |
| DRC: Mikalayi | *Anopheles moucheti* | DRC-MIK1 | W+ | MH598424 |
| DRC: Kinshasa | *Anopheles gambiae* s.s. | DRC-KIN1 | W+ | MH598425 |
| DRC: Mikalayi | *Anopheles gambiae* s.s. | DRC-MIK2 | W+ | MH598426 |
| DRC: Kalemie | *Anopheles gambiae* s.s. | DRC-KAL1 | W+ | MH598427 |
| DRC: Kalemie | *Anopheles arabiensis* | DRC-KAL2 | W+ | MH598428 |
| DRC: Katana | *Anopheles gambiae* s.s. | DRC-KAT3 | W- | MH598429 |
| Ghana: Dogo | *Anopheles coluzzii* | GHA-DOG1 | W+ | MH598430 |
| Ghana: Dogo | *Anopheles coluzzii* | GHA-DOG2 | W+ | MH598431 |
| Ghana: Dogo | *Anopheles coluzzii* | GHA-DOG3 | W+ | MH598432 |
| Ghana: Dogo | *Anopheles coluzzii* | GHA-DOG4 | W- | MH598433 |
| Ghana: Dogo | *Anopheles coluzzii* | GHA-DOG5 | W- | MH598434 |
| Ghana: Dogo | *Anopheles melas* | GHA-DOG6 | W- | MH598435 |
| Uganda: Butemba | *Anopheles gambiae* s.s. | UGA-BUT1 | W- | MH598436 |
| Uganda: Butemba | *Anopheles gambiae* s.s. | UGA-BUT2 | W- | MH598437 |
| Uganda: Butemba | *Anopheles gambiae* s.s. | UGA-BUT3 | W- | MH598438 |
| Uganda: Butemba | *Anopheles gambiae* s.s. | UGA-BUT4 | W- | MH598439 |
| Uganda: Butemba | *Anopheles arabiensis* | UGA-BUT5 | W- | MH598440 |
| Madagascar: Antafia | *Anopheles gambiae* s.s. | MAD-ANT1 | W- | MH598441 |
| Madagascar: Antafia | *Anopheles gambiae* s.s. | MAD-ANT2 | W- | MH598442 |
| Madagascar: Antafia | *Anopheles gambiae* s.s. | MAD-ANT3 | W- | MH598443 |
| Madagascar: Ambomiharina | *Anopheles rufipes* | MAD-AMB1 | W- | MH598444 |
| Madagascar: Ambomiharina | *Anopheles rufipes* | MAD-AMB2 | W- | MH598445 |

**Supplementary Table 2.** *Wolbachia* 16S and wsp GenBank accession numbers

| Sample ID | Strain | 16S | wsp |
| --- | --- | --- | --- |
| DRC-LWI1 | wAnsA | MH605275 | MH605281 |
| DRC-LWI2 | wAnsA | MH605276 | MH605282 |
| DRC-LWI3 | wAnsA | MH605277 | MH605283 |
| DRC-KAT1 | wAnsA | - | MH605284 |
| DRC-KAT2 | wAnsA(2) | - | MH605285 |
| DRC-MIK1 | wAnM | MH605278 | - |
| GHA-DOG1 | wAnga-Ghana | MH605279 | - |
| GHA-DOG2 | wAnga-Ghana | MH605280 | - |

**Supplementary Table 3.**
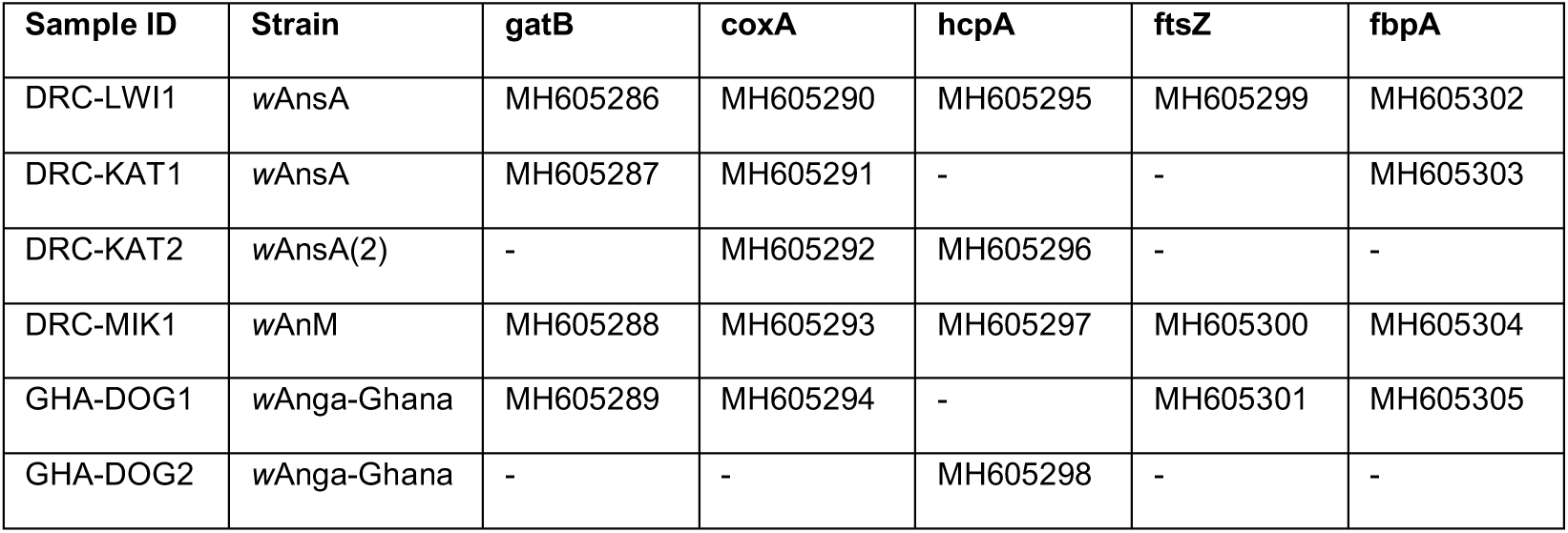
*Wolbachia* MLST gene GenBank accession numbers

